# Node Features of Chromosome Structure Network and Their Connections to Genome Annotation

**DOI:** 10.1101/2023.12.29.573476

**Authors:** Yingjie Xu, Priyojit Das, Rachel P. McCord, Tongye Shen

## Abstract

The 3D conformations of chromosomes can encode biological significance, and its implication is being increasingly appreciated recently. Certain chromosome structural features, such as A/B compartmentalization, are frequently extracted from pairwise contact information (physical association between different regions of the genome) and compared with linear annotations of the genome, such as histone modifications and lamina association. Here, we investigate how additional properties of chromosome structure can be deduced using the abstract graph representation of the contact heatmap, and how network properties can have a better connection with some of these biological annotations. We constructed chromosome structure networks (CSNs) from bulk Hi-C data and calculated a set of site-resolved (node-based) network properties of these CSNs. We found these network properties are useful for characterizing chromosome structure features. We examined the ability of network properties in differentiating several scenarios, such as haploid vs diploid cells, partially inverted nuclei vs conventional architecture, and structural changes during cell development. We also examined the connection between network properties and a series of other linear annotations, such as histone modifications and chromatin states including poised promoter and enhancer labels. We found that semi-local network properties are more capable of characterizing genome annotations than diffusive or ultra-local node features. For example, local square clustering coefficient can be a strong classifier of lamina-associated domains (LADs), whereas a path-based network property, closeness centrality, does not vary concordantly with LAD status. We demonstrated that network properties can be useful for discerning large-scale chromosome structures that emerge in different biological situations.

**TOC Figure:** 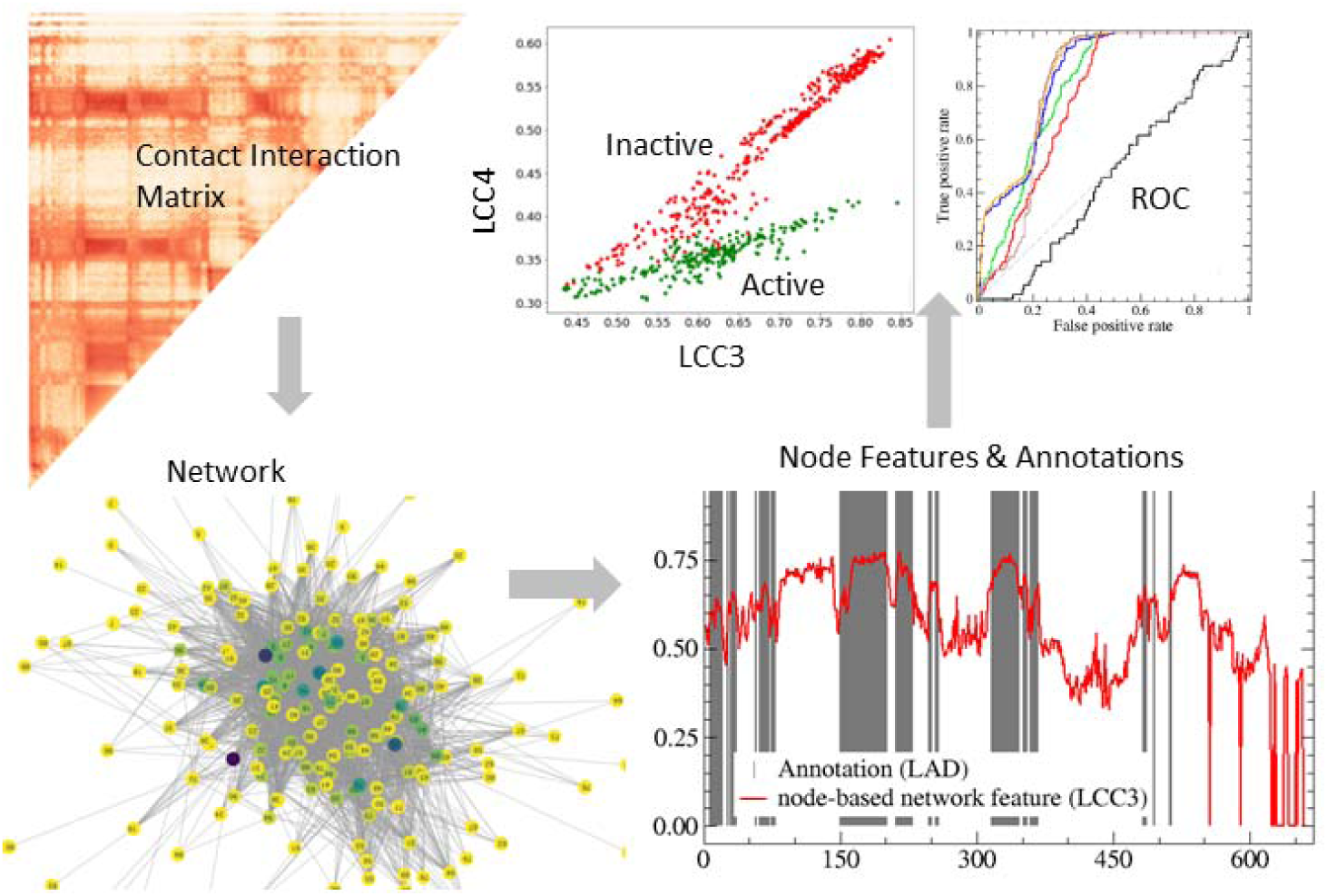

## I Introduction

The chromosomes of eukaryotes are arranged in a multi-scale and complex fold inside the nucleus which can be important for their biological functions (1). The long-range spatial structures of chromosomes can contribute to the regulation of gene expression by fostering enhancer-promoter contacts locally through loop extrusion or over longer distances (and between chromosomes) through spatial compartmentalization (2,3). Variation in chromosome structure can both be an important source of functional diversity and a cause in certain pathological traits (4–6). Chromosome structure capture methods have enabled the study of chromosome organization at the genome scale (7,8). Hi-C is a high-throughput chromosome 3D structure capture method that can reveal population-based (bulk) chromosome structures expressed by pairwise contact interactions. A typical Hi-C analysis uses a two-dimensional contact map that indicates the frequency of contacts between all possible pairs of genomic positions (genomic bins), as illustrated in Fig. 1. However, this two-dimensional matrix might not be intuitive to grasp, and it is difficult to visualize the correlation between the spatial structure and to make connection to biological annotations, which are often associated with single genome positions. Therefore, extracting one-dimensional (1D) indicators from two-dimensional (2D) contact map is a useful way to characterize chromosome structure features, compare them to other linear genomic features, and discern subtle differences between cell types or upon perturbations.

**Figure 1.**
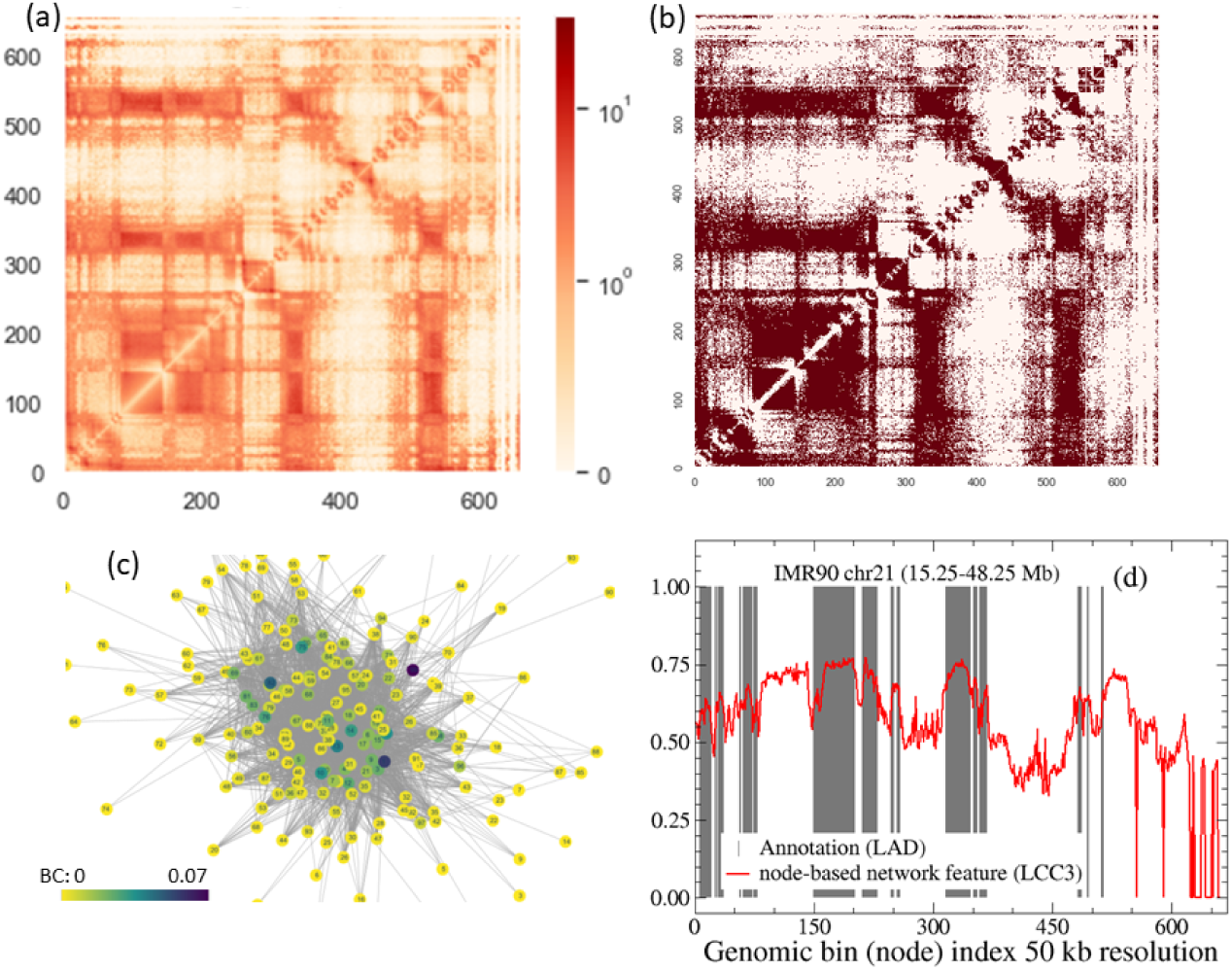
(a) Hi-C contact map of IMR90 chr21 (region 15.25–48.25 Mb) at 50 kb resolution (b) The corresponding network adjacency matrix representation with a cutoff of 40% coverage (c) The corresponding force-directed network graph representation of panel (b). (d) Network feature (LCC3) output (blue) and LAD annotation (gray) as functions of node (genomic bin) index.

One commonly seen 1D indicator is A/B compartment strength (which is often further discretized to a binary A/B classification) for each genomic bin position. A/B compartment strength is obtained from the principal component analysis (PCA) of bulk Hi-C-derived contact matrices. There are other ways of rendering 2D contact matrices to 1D genomic bin-based information, from the insulation score or the directionality index to define TAD boundaries (9,10), GAD score (11), to distal-to-local ratio (DLR) and interchromosomal fraction (ICF) (12) which have been used to measure dynamic properties such as Loss of Structure (LOS) after perturbation (13). Here, we explore a set of systematic rendering methods using abstract graph representation and network analysis. We are interested in both evaluating how these mathematical concepts relate to each other and the connection between the underlying genome structure and the biological linear annotation.

Discretizing spatial connection (contact interaction strength) and constructing graph representations of structural information of biomolecules and larger biological structures facilitates the application of a family of node-based network properties (Fig. 2a), such as centralities and clustering coefficients, to understand biological functions. At a relatively small size scale, network analysis is applied to study chemical structures (14) and internal interaction of macromolecules such as proteins. Protein structure networks have been extensively discussed and shown to be useful for studying protein structure, dynamics, (15–22). Beyond single molecules, abstract graph theories have been used to describe the structure and reaction dynamics of molecular networks of chemicals, from the oxidation of oil paintings to the sol-gel phase transition of polymers (23,24). It is interesting to examine how such abstract graph representations of structures can be connected to biological function and biophysical aspects of complex and heterogenous chromosome structures.

**Figure 2.**
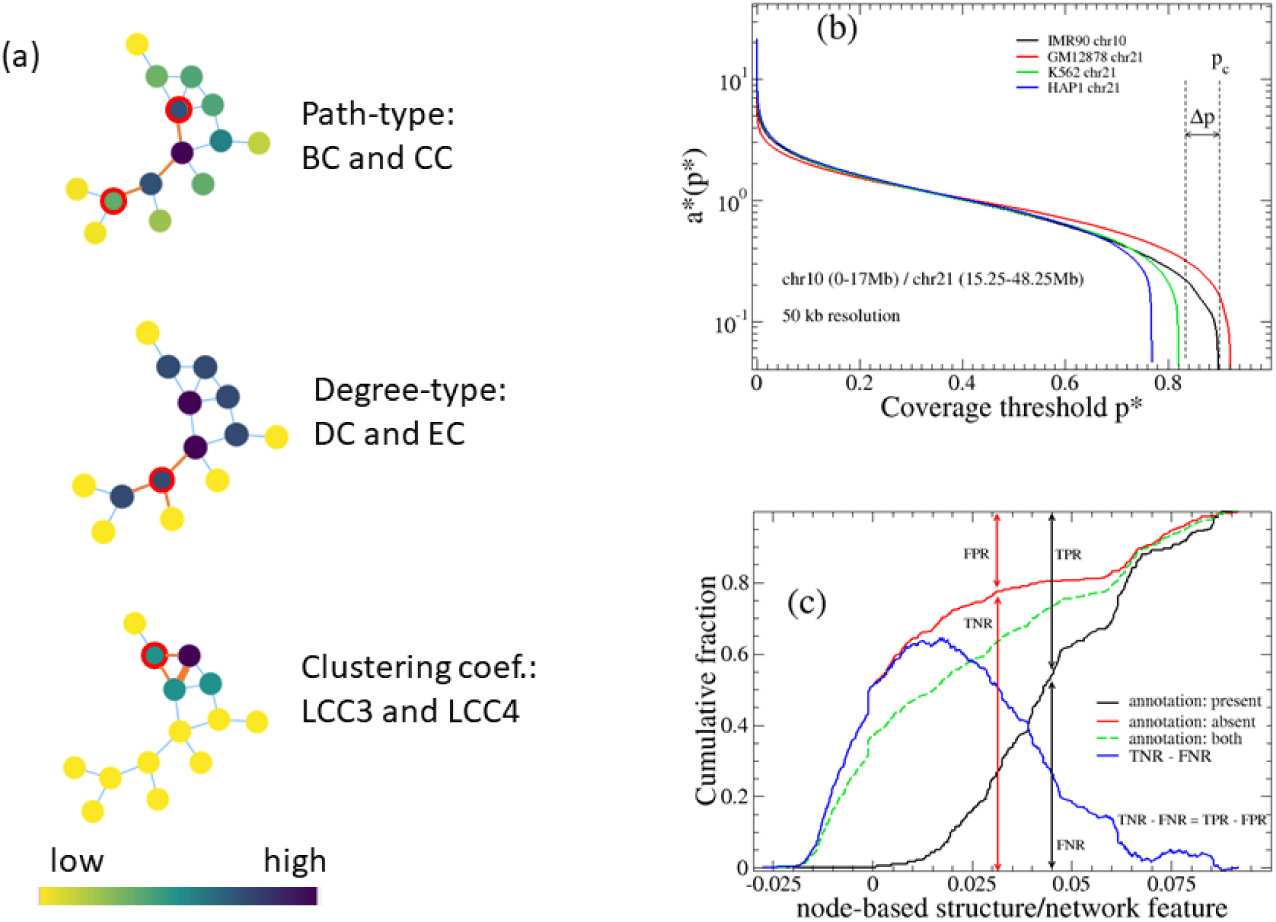
(a) Pedagogical graphs are shown as illustrations of several network properties. The relative value of network property for each node is color labeled. Both BC and CC are path-oriented measurements. BC counts the paths that go through a target node while CC focuses on the paths terminating at a target node. In contrast, DC and EC focus on the number of neighbors of the target node. LCC indicates the number of closed local paths a node has. (b) The coverage (normalized rank of contact) *p*\* shown as a function of contact strength *a*\*. Δp indicates the coverage margin. (c) Cumulative fraction curves for a label being present (black), absent (red), and reference (green). ROC is further defined based on associated concepts TPR, TNR, FPR, and FNR.

Previously, the applications of network analysis for genome biology have been largely centered on molecular interaction networks and co-expression networks (25–27) whereas few studies focused on the structure networks and the node-based network properties of chromosomes. In the current study, we explore the node-based features that reflect various aspects of interaction between a given chromosome region (a genomic bin) with its spatial neighbors. Some of the node features are utterly local, such as the degree centrality, while others are quite global, such as closeness and betweenness centralities (20,28). It is interesting to examine whether specific features of networks reflect strongly a specific biological feature and to what extent network analysis can reveal interesting structure-function relationship within a chromosome and between individual chromosomes due to cell type variation and environmental influences.

One example of the influence of the chromosome territory environment on chromosome structure is found when one compares wild type (WT) thymocytes, its lamin B receptor (LBR) mutant and rod cells. Interphase nuclei of WT thymocytes represent a conventional architecture: heterochromatin regions are primarily located at the nuclear periphery, whereas euchromatin (typically, regions that are gene rich and largely active) resides in the nuclear interior. FISH experiments (29) have revealed that the structure is inverted in rod photoreceptor cells of nocturnal mammals, and likewise partially inverted in LBR -/-mutant thymocytes (30). The previous direct comparison of Hi-C data between such inverted cells and the normal counterpart found straightforward changes between rod cells and WT thymocyte, but the difference between LBR mutant and WT thymocyte is subtle and the A/B compartment representation was unchanged despite the partial inversion. Here, we ask whether distinct features can be extracted by network properties and how a network viewpoint enhances our understanding of this type of biological comparison.

Another application of graph representation is the influence of cell differentiation on chromosome structure. Here, we inspect how chromosome structural features are altered between progenitor cells and differentiated cells using a set of blood cell types (31). Other examples include how chromosome structure in a haploid cell is different from that in a diploid cell, the differences between a cancer cell type and its normal counterpart, and how the nuclear lamina-chromosome interaction is reflected in the chromosome structure network (CSN).

## II Methods and systems

### A. Contact network construction from chromosome structure information

Abstract graph is a useful way of representing structural components and their relationship. For each graph (network), two types of elements, vertices and edges, are present. The basic structural unit is termed a vertex, which is also called a node in computational sciences or a site in physical sciences; whereas an edge that connects two vertices is termed a link in computational sciences or a bond in physical sciences. We will use these terms interchangeably. In this case, each node is a chromosome region (genomic bin) of size 50 kb unless specified otherwise, whereas each link indicates two such regions are close spatially and their association (contact interaction) can be detected in Hi-C experiments.

The bulk Hi-C data can be represented as a contact matrix A, where *a_jk_* indicates the number of contacts recorded between regions *j* and *k*, assuming that each region is indexed from 1 to N. There are total N nodes in the system. Though self-edges are typically included in a graph representation and are used for downstream analyses, those contacts (diagonal elements) are not considered for the current analysis. Essentially, we would introduce a cutoff value to discretize the values of *a_jk_* to either 1 when *a_jk_* ≥ *a*\* or 0 otherwise. Here 1 indicates a link (edge) is formed between nodes j and k. Such discretized contact matrix is termed the adjacency matrix of a network. The discretization and construction of graph representation helps us simplify the relationship between nodes. However, one needs to be careful when choosing a threshold. There are multiple ways of defining the threshold and each has a different emphasis. For example, one uses a random polymer model as a reference and considers link formed if the contact value is more than the reference value, such as FitHiC method (32,33). Such a method might provide different answers depending on data resolution and it may convert to an overly saturated network in practical cases, especially when the resolution is low and bin size is large.

In this work, we select threshold by using the median value of nonzero contact strengths and thus maximizing Shannon Entropy of edge formation. Essentially half of the N(N-1)/2 pair of nodes is connected. Note that this threshold selection is sensitive to the data quality, i.e., the normalized number of nonzero *a_jk_*. A properly selected threshold value *a*\* sensitively depends on experimental conditions such as sampling size. Since the meaning of cutoff value *a*\* is not intuitive for selecting links, instead, we define a concept termed link saturation (coverage) *p*\*, a quantitative comparison with the complete network, where all edges are formed. With a running cutoff, we can plot the function *a*\*(*p*\*) as a curve to connect these two concepts. As shown in Fig. 2b, the j*th* ranked contact with strength *a_j_* contributes to a point 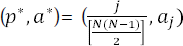 on the curve *a*\*(*p*\*) . When *a*\* is 0, all links are formed and one obtains a complete graph where there are total *N*(*N*-1)/2 links in the graph. With increasing *a*\*, some of the links are removed. Parameter *p*\* can be defined as the ratio between the number of total links of a graph and the link number of a complete graph. We can use a cutoff link saturation (*p*\*) to specify *a*\* and further define the contact network, which means only the largest contacts (*p*\* ×100 %) were considered formed. The proper cutoff value ensures that network properties maintain structure information while at the same time, the cutoff filters out other contacts deemed insignificant.

Note that practically *p*\* may not always be set a high value that is close to 1, as it is affected by how many zero elements the contact matrix has which in turn is affected by several factors. For one, certain regions of the chromosome are highly repetitive and cannot be easily resolved. As a result, when there are only total N_c_ nonzero values, we can define 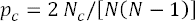. Since p_c_ represents the maximum link saturation at a contact threshold of zero, clearly *p*\* should be less than *p_c_*. Other factors that influence the sampling of a contact map include crosslinking duration and sequence depth. These factors create another layer of uncertainty. Practically we can define the number of elements *a_jk_* that only have a single “hit”, *N*_1_, and a safty margin parameter 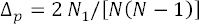. Together we have 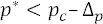. Ideally one could choose 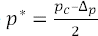. Note that the raw (discrete) number of contact counts is only used for selecting proper threshold *p*\*, whereas *a*\* requires iterative correction (ICE balancing (34)) to adjust biases in Hi-C data collection before one can render a network representation. Here *N*_1_ and *N_c_* indicate the quality of the sampling. For example, for the whole chr21 of IMR90 dataset (35) at 250 kb resolution, *p_c_*= 53.1% and Δ_p_= 1.2%. Thus, one should not choose *p*\* more than 50% in such a case. In Fig. 2b, we plot threshold value *a*\*vs cutoff link saturation *p*\* for chr10 (region 0-17Mb). One can see that a small (tolerant) *a*\* threshold will include all the nonzero elements of the contact matrix and results a near maximum coverage *p*\*Note that *p*\*does not necessarily achieve 100% as not all genome bin interactions are detected, especially at higher resolutions. For the case of IMR90 chr10, *p_c_* = 89.9% and the coverage margin parameter Δ_p_ = 6.6%. Ideally one should choose *p*\* at the flattest region of the *a*\*(*p*\*) curve since such cutoff will make the selection least sensitive to the choice of a*. According to the results, it is better to choose a cutoff of 40% under 250 kb resolution. Practically, we use *p*\* =40% unless specified otherwise.

### B. Network properties

Once the discretization procedure of contact matrix is achieved and abstract graph is constructed (Fig. 1a-c), we can further calculate site-based network properties. For a given network, there are a range of node-based network properties one can construct, from properties that are quite local and reflect only the connectivity of how a genomic bin with its linked neighbors to those global and collective in nature. Here, we mainly study centrality properties, clustering properties, and a hybrid of centrality and clustering properties. As a comparative overview, the two centralities (closeness centrality CC and betweenness centrality BC) that are defined from paths are quite global, while two eigensystem based properties, eigenvector centrality (EC) and A/B strength (MI-PCA) are semi-local, followed by local clustering coefficients (LCCs) and finally degree centrality (DC) is the most local. The order (from global to local node features) is: CC and BC, EC, …, LCCN, LCC(N-1), …, LCC4, LCC3, and DC.

Roughly speaking, betweenness centrality (BC) measures how often a node appears on shortest paths between two nodes while closeness centrality (CC) focuses on nodes being ends of a shortest path (Fig. 2a). Both properties are path-based parameters and emphasize the global features of the network. In contrast, degree centrality (DC) and eigenvector centrality (EC) measure local properties. Both DC and EC count the number of neighbors of a node, but the main difference is that EC considers a self-consistent weight assigned to each node and it is more global than DC. Highly connected nodes weigh more than the low ones. The local a link exists between *i* and *j*, 0 otherwise. clustering coefficient (LCC) is another way of associating network features to node-based values. LCC is a way of indicating how dense the links are around a node, and in comparison, it is more local than some of the centralities. Specifically, the LCC value for node *i* is given by 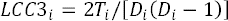 (28). The term *T*j represents the count of triangles that include vertex *i* and 2/[*D_i_*(*D_i_*-1)], the total possible triangles given the degree of ith node, *D_i_*, which defines the number of neighbors of vertex v. Using adjacency matrix, A, one can generalize LCC3 and describe completed neighboring squares for ith node (36,37), 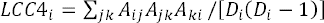 where 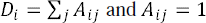 when a link exists between *i* and *j*, 0 otherwise. As we will demonstrate below, LCC4 (LCC-even, in general) is largely independent from LCC3 (LCC-odd) and especially useful for studying certain types of networks such as bipartite networks (38). Both provide distinct perspectives of network features. Besides the loop-based generalization we adopted, there are some other expansions of definitions, such as clique-based LCCs (39) that can provide an additional insight on the architecture of networks in general.

### C. Statistical analysis

Although site-based contact PCA is not deduced from properties of a network, it is a popular way of reducing 2D contact map information to site-based values (40). For MI-PCA and downstream discretization of A/B compartment analysis, each element *x_j_* of the top eigenvector (one with the largest eigenvalue) is associated with each genomic bin *j* and the A/B compartment is assigned to each bin depending on the sign of the eigenvector element at that bin. As one group of elements is often associated with gene rich, while the other gene poor, one can assign the signs so that *j* belongs to the A compartment when *x_j_* >0 and belong to B when *x_j_*< 0. Absolute value |*x_j_*| also is used to symbolize the strength of the compartment.

We would like to quantitatively compare local, genomic bin-associated network properties and linear annotations of linear genome information which are often expressed as discrete (even binary) states. The conventional scatter plot and correlation coefficients may not work in such cases. Instead, we report the area under the cumulative fraction curves (CFCs) which are essentially an integration of the raw scatter plot function (e.g., the binary state on y-axis is being converted to “stairs”).

For each pair of network properties and binary annotation information, one can construct three cumulative fraction curves, CF_P_ (black), CF_N_ (red), and CF_R_ (green dashes), as illustrated in Fig. 2c. Each genomic bin position has a network value, e.g., LCC3 and a binary annotation, e.g., being LAD or not LAD. For CF_P_, the cumulative fraction curve examines the positive response and increases by a fraction of *d_y_* = 1/*N_p_* at position x = LCC3_i_, when i*th* genomic bin is LAD and *d_y_* = 0 when it is not LAD. Here *N_p_* is the total number of nodes with LAD status. Vice versa, CF_N_ examines the negative response and increase when it is not LAD by *d_y_* = 1/*N_n_*. And *N_n_* is the total number of nodes that is not LAD. The reference curve CF_R_ increases by *d_y_* = 1/(*N_n_*+*N_p_*) at LCC3_i_ regardless of LAD status. Now when we make a hypothesis that larger values of LCC3 are associated LAD status, the value of CF_P_ is the false negative rate (FNR) while 1-FNR is true positive rate (TPR). Meanwhile, one can obtain true negative rate (TNR) and false positive rate (FPR) from the CF_N_ curve. We can further define TNR-FNR (=TRP-FRP) and find an optimal threshold value to classify these two states (LAD or not) from LCC3 values. Note that the positive likelihood ratio LR+ =TNR/FNP is a similar way of discerning these two curves. Further, the widely applied receiver operating characteristic (ROC) curve is plotted (x, y) = (FPR, TPR). The area under the ROC curve (AUC) is a value between 0 and 1 and a larger value implies a stronger performance measurement for the classification of an annotation from a given network property.

### D. Systems and data

1. Hi-C data: All structural data of chromosomes used in this study came from bulk Hi-C data. In this study, we have used publicly available (www.ncbi.nlm.nih.gov/geo) Hi-C data from the following human cell types: lymphoblast GM12878 (35), fibroblast IMR90 (35), leukemia cell line K562 (35), HAP1 – a haploid cell derived from K562 (41), and data of different blood cell types: erythroblasts, neutrophil and megakaryocyte with permission from the PCHI-C Consortium (31). Additionally, we also studied Hi-C data of mouse cell types: WT vs. LBR mutant thymocytes for comparing the effects of overall nucleus architecture on network properties (30). Hi-C raw data is expressed as a contact frequency (number of “hits”) between regions of the genome at a specific resolution (genomic bin). With proper procedures one can describe the information as a contact matrix. Unless otherwise specified, all Hi-C data came through MAPQGE30-filtering and was further normalized by ICE balancing (34). Furthermore, we apply a sequence distance normalization on contacts diagonal, i.e., 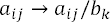 with 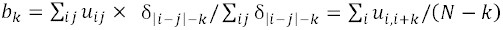, where Kronecker delta selects *k* = |*i*-*j*|, as stated in Ref. (40). Previously, we pointed out that unlike ICE balancing which is a necessary site-based correction, both distance-normalized and untreated versions can be useful, where the former one is focused on interactions that occur more often than expected in a random polymer, and therefore perhaps are mediated by specific biological mechanisms, while the latter represents how often two regions actually come in contact in a nucleus, which has implications for biochemical reactions (40). Since here we focus on study structure patterns, we use distance normalization for all data presented.

We focus on intra-chromosomal interactions in this study, and chromosome 21 is examined by default. Since a part of the chr21 is poorly covered by Hi-C experiment due to repetitive nature, we used the 15– 48 Mb region. To generalize our conclusion, we also examined another (larger) chromosome, chr10 (∼ 0– 17 Mb) for a comparison.

There are many interesting linear annotation data on chromosomes. There are several important aspects, from chromosome histone modification marks, lamina-associated domains (LADs), gene expression, to derived information such as chromosome sub-compartments and chromatin states. We focus on examining chromatin state, LADs, and compartment subtype in this work. We choose these biological properties since they are closely associated with structural properties of chromosomes and thus may manifest as network properties or biologically significant.

2. Lamina-Associated Domain: Lamin is a nuclear inner membrane protein that is critical for various biological processes within the nucleus, such as chromatin organization, DNA repair, and gene expression. In mammals, both A- and B-type of lamins interact with hundreds of large chromatin domains known as lamin-associated domains (LADs) (42). The data of chromatin-lamin interaction came from Ref. (42). Specifically, LADs are spatially located at the periphery of the nucleus, adhering to the nuclear membrane. LADs might be closely tied to geometrical properties of chromosome structure that are therefore reflected in a graph representation. LADs also change their spatial localization in response to cell type specific gene expression during differentiation and development (43). For this study, LAD data came from Tig3 cells. However, there is no Hi-C data for Tig3 cells presently available. Since Tig3 cells are fibroblasts similar to IMR90, we make a comparison of Tig3 LADs with Hi-C results of IMR90 below.

3. Chromatin state: Chromatin states (CSs) map epigenomic marks such as histone modifications, histone variants, open chromatin regions, and associated marks to their likely functional roles in gene regulation (44). Each region of the chromosome is assigned to only one of the 15 CS states according to their combination of epigenetic marks: CS1=Active Promoter, CS2=Weak Promoter, CS3=Poised Promoter, CS4=Strong Enhancer 1, CS5 =Strong Enhancer 2, CS6 =Weak Enhancer 1, CS7=Weak Enhancer 2, CS8=Insulator, CS9 = Transcription Transition, CS10= Transcription Elongation, CS11=Weak Transcription, CS12=Repressed, CS13=Heterochromatin/low signal, CS14=Repetitive/ Copy Number Variation, CS15=Repetitive/ Copy Number Variation. We used CSs of GM12878 in this study.

Note that the annotation state of a genome sequence, such as chromatin state, exists at a different (and often higher) resolution than that of the network nodes. In the current case, CS data has a much higher resolution of 200 bp. When we examine cumulative fractions and make comparisons between chromatin states and network properties, we operate at the lower resolution (e.g., 50 kb) of the two datasets and directly assign the annotation state of the exact genome position of the node (the lower bound of the corresponding genome bin), not a more sophisticated method such as the majority rule of a running window centered on the node position. Our straightforward definition works well for correlation calculations. However, when aiming to depict the statistical characteristics of network properties for a particular chromatin state using a distribution (violin plot), say that for CS3 (poised promoter), we faced the problem that there are few nodes assigned to this label, because this annotation is typically very small and does not get assigned to the larger genomic bin node. To resolve this issue, we practically operate at the higher resolution (0.2 kb) mode and interpolate by assigning the same network property value to all bins that fall within each 50 kb range (one structure node). Thus, for the violin plots alone, multiple chromatin state labels can be assigned to the same node and quantitative weights are assigned.

4. Subcompartments: In addition to the binary classification of A/B compartment (largely correlated with euchromatin vs heterochromatin) (45), there are further studies that classify regions into multiple compartment subtypes (sub-compartments) using various information, from interchromosomal contacts, strength of PCA eigenvectors, gene expression, and epigenetic information (35,46–49). Here we compared our network properties with the Rao et al. sub-compartment definition, which extends the A/B definition to six states, A2, B1, B2, B3 and B4 (45).

5. Mouse Interphase nuclei: Another application is studying the effect of overall genome architecture and chromosome territory positioning on their network features. Here we compare different nucleus architectures. Interphase nuclei have a conserved (conventional) architecture: heterochromatin typically occupies the nuclear periphery, whereas euchromatin resides in the nuclear interior. Meanwhile, rod photoreceptor cells of nocturnal mammals have an inverted architecture, which transforms these nuclei into microlenses and facilitates a reduction in photon loss in the retina (30). Furthermore, a partially inverted architecture was observed when lamin B receptor is deleted from thymocytes. Compared to WT thymocytes, these LBR mutant cells cannot anchor lamin associated domains onto the nuclear membrane. Though it is relatively simple to discern the differences in Hi-C maps from WT thymocytes (conventional) and rod cells (inverted) because these are quite different cell types, it is difficult to distinguish between LBR -/- (partially inverted) and WT (conventional) architecture using Hi-C contact maps directly. Therefore, we examine whether network properties can distinguish these two nuclear organizations based on their subtle changes in Hi-C data. We used a gene dense mouse chr11 for this study. The original data resolution is 20 kb, and we use 200 kb for better sampling in the network analysis.

## III Results and discussion

### A. Effects of coverage and resolution on chromosome structure network properties

We first report how resolution and threshold might affect the results using chr21 of IMR90 (region 15.25– 48.25 Mb) as an example. By varying coverage parameter *p*\*, we found that a higher cutoff value such as 80% results in constant high values of the node-based network property LCC3, as the nodes of the CSN are almost fully connected. On the contrary, if the threshold *p*\* is as low as 20%, the resulting curve has larger variation with some nodes being zero, as CSN has fewer links and is being broken into disconnected smaller subnetworks (Fig. 3a). These results suggest a *p*\* threshold of 40%, the same as chosen based on observations from Fig. 2b, is also appropriate to yield network properties that vary meaningfully across the chromosome.

**Figure 3.**
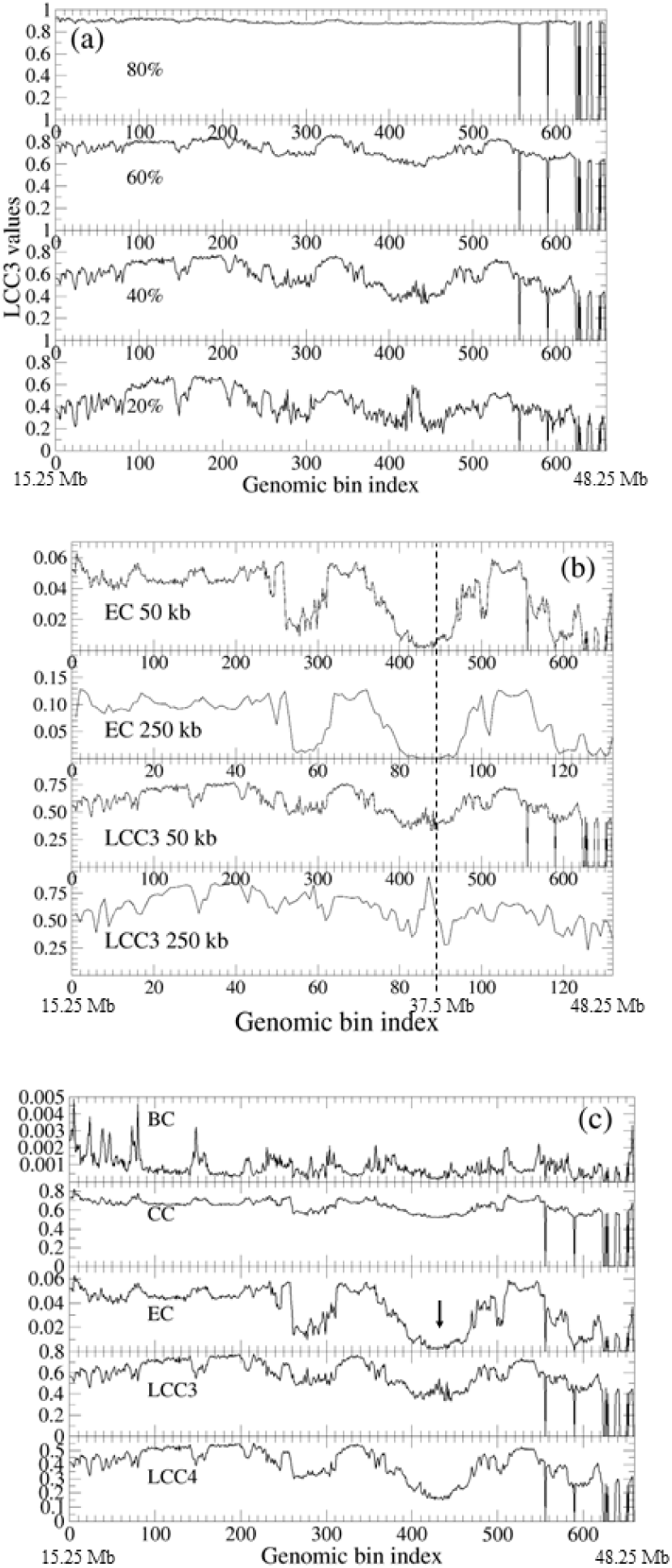
(a) LCC3 value as a function of node (genomic bin) index, under different coverage thresholds *p*\* (max, 80%, 60%, 40%, 20%) at 50 kb resolution for IMR90, chr21:15.25–48.25 Mb. (b) IMR90 chromosome 21 (same region as above) 40% *p*\* cutoff EC and LCC3 values at different resolution (50 kb vs 250 kb bin sizes). (c) Comparison of different network properties including BC, CC, EC, LCC3 and LCC4 across the same IMR90 chr21 region as in a and b. Arrow indicates a notable region of divergent values between network properties at a given genomic location.

Resolution also has an impact on network properties, as shown in Fig. 3b. For example, there is a potentially spurious high peak for LCC3 values near 37.5 Mb (bin index 89) while the peak disappears if the resolution is 50 kb (bin index 445), when the genomic bin size is decreased from 250 kb to 50 kb. We observe that the resolution has a relatively small effect on EC, which indicates eigenvector centrality is less sensitive to resolution selection since EC is more of a semi-global property compared to local properties such as LCC3. Generally, a smaller bin size provides a high-resolution description only if there are sufficient samples in each bin.

We next compare the similarities and differences in the quantified network properties, LCC3, EC, BC, CC, and LCC4, using the same parameters (250 kb and *p*\*= 40%) (Fig. 3c). BC exhibits the most frequent variations, while the other four properties show changes more gradually across the genomic region. Overall, BC values are nearly constantly low while CC values are constantly high. EC and LCC3 have a wider range of variation. But their peaks and valleys are not correlated, which indicates different network properties reflect distinct features of the CSN. Using several different network properties may provide a more comprehensive understanding of chromosome structure. For example, at positions around 35,000∼37,500 kb (arrow, Fig. 3c), the EC is almost zero, the LCC4 value is around 0.3, and the LCC3 value is high. The low CC and EC suggests that the region has few neighbors, while the contrasting LCC3 and LCC4 values indicates that there is a short-distance ring structure in the area. Thus, this region is like an isolated island, likely to form links among themselves rather than with others.

The network properties we quantify provide different measurements of the structure features, but previous research pointed out possible correlations between derivative network properties, such as BC vs DC x (1-LCC3) (50). We directly compared the correlation between different network properties and between network properties and A/B compartment strength in Supplementary Material (SM) Fig. S1. Among all the network properties studied, most show poor correlation. Only a few pairs of properties show correlation in restricted regions. For example, LCC4 and A/B compartment strength show correlation at high value regions whereas EC and A/B correspond at low value regions. We found only one strongly correlated pair: EC and DC x LCC3. Additionally, as shown in SM Fig. S2, a derived property, LCC4/LCC3 ratio, which is a relative comparison of the strength of short-range connectivity (nearest neighbors) vs. relatively global connections (neighbors of nearest neighbors), is correlated to A/B compartment classification. Thus, nodes with high LCC4/LCC3 ratios are likely a part of the inactive region (B compartment) and vice versa, which further demonstrates that graph properties can be a useful characterization of biologically related structure features.

### B. Network properties display distinct structural features of chromosomes for different cell types

There are various epigenetic and genetic differences between chromosomes of different cell types. We next asked whether network properties and CSN can discern these chromosome structural feature differences exhibited in different cell types. We first compared the CSN of chr21 between a normal cell type (GM12878, non-cancerous lymphoblast) and a cancer counterpart (K562; leukemia). As shown in Fig. 4a, the overall fluctuation of LCC3 and LCC4 values of K562 is much more subdued than the corresponding features of normal cells (GM12878). This flatter pattern, which does not vary significantly between sequence neighbors, suggests that many nodes on a chromosome have a similar (and even) degree of connectivity with other nodes in the network. The loss of network feature in bulk cancer cell data may reflect the loss of functional organization of the chromosome. Thus, unlike normal cells which have highly organized chromosome structure, the degree of chromosome organization in cancer cell K562 is weakened. Although most of the remaining peak and valley positions of K562 are similar to that of GM12878 (especially for EC), there are exceptions. For example, K562 is missing the peak at about 37 Mb for LCC3, potentially a site of more dramatic chromosome structural difference.

**Figure 4.**
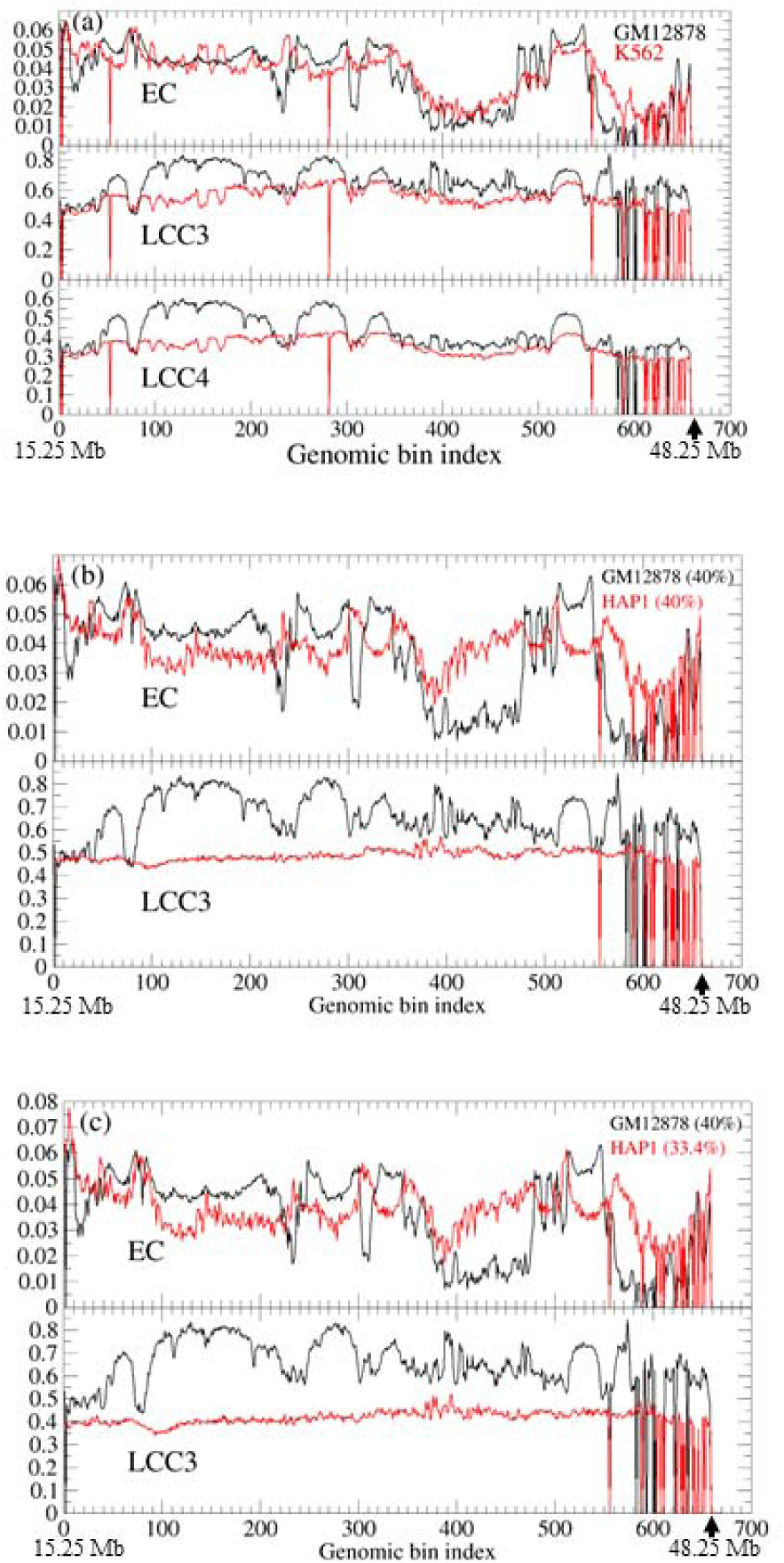
Network properties (EC and LCC3/4) as functions of network node index for cancer cell type K562 are shown in (a) and for haploid cell HAP1 in (b). The corresponding results for lymphoblast GM12878 are shown as a comparison. Data of region 15.25–48.25 Mb for chr21 (50 kb resolution) is used. Constant coverage of 40% is used. (c) Same comparison as (b) but HAP1 coverage is set to 33.4%, so that *p*\*/*p_c_* is kept constant.

In addition to changes in chromosomal structure in cancer, we also examined the impact of interactions between homologous chromosomes on genome structure. In this case, haploid cells HAP1 vs. diploid cells GM12878 were compared. HAP1 has only a single copy of each chromosome except chromosome 15 (the second copy of chr15 is only an incomplete fragment). Therefore, chromosomes of haploid cells may lose some of the interchromosomal interactions of the corresponding diploid cells. We constructed contact networks for HAP1 using the same cutoff (*p*\* = 40%) and displayed the EC and LCC3 results in Fig. 4b. It is worth mentioning that there is a noticeable difference in the coverage of Hi-C sequencing data, where GM12878 has a higher *p_c_* than HAP1 (*p_c_* =92.5% vs 77.2%).

When using the same cutoff *p*\*, the system with lower *p_c_* has fewer dispensable edges to choose and thus leads to a structure network that is more mean-field like, an evenly connected global network. Since *p_c_* (data coverage) influences the effect of *p*\*, we also used an alternative definition, a scaled *p*\* to compare different systems. Specifically, instead of using a constant *p*\*, we use different *p*\* for different systems while keep 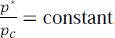. For this case, we make sure that 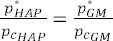. As shown in Fig. 4c, we set *p*\*=33.4% to construct the network of HAP1, to ensure 33.4%/77.2% = 40%/92.5%. When we compare Fig. 4b and 4c, the results are not sensitive to the selection of *p*\*, i.e., lowering the cutoff value of HAP1 cells, did not significantly affect LCC3 and EC values. Similar to the cancer cell K562, from which HAP1 is derived, the LCC3 values are almost constant, and each position has a similar number of random contacts with its surrounding network. Subsequent tests showed that even if *p*\* is lowered to 20%, the LCC3 value of HAP1 would still be largely constant with slightly more random fluctuation (SM Fig. S3). Thus, this phenomenon is robust and not an effect of *p*\*, which suggests the lack of well-defined chromosome structure features in HAP1. Since HAP1 is both haploid cell and originates from cancer cell line, it is unclear whether one or both factors contribute to the phenomenon. However, upon examining the EC values, we observe that HAP1 further weakens some of the organizational structure (for example, in the 19.5–24 Mb region, SM Fig. S3) compared with K562, which suggests that some of the structural changes might be caused by the haploid nature of the cells.

Rod cells in mice have a unique inverted nuclear structure. It is difficult to see the difference between the “inverted” and “non-inverted” structures from the Hi-C contact map directly. We analyzed the differences between WT type and LBR mutation using the chromosome structure networks. As shown in Fig. 5, on the whole, BC, CC and EC show a similar distribution pattern of high values at both ends and low in the middle. On the contrary, The LCC value is higher in the middle than at both ends. This suggests that the nodes in the central region of mouse chr11 have fewer neighbors but are more locally connected whereas the nodes at the ends are more urban. A scenario of long-distance enhancer-promoter interaction at the ends and highly correlated regulation complexes in the middle can be a realization of such CSNs. Compared with the wild type, the value of network properties fluctuated less in the LBR mutant. And the LCC values of LBR mutant are lower than that of wild type while BC and CC do not change much, which suggests that the LBR mutation has weaker short-distance contacts and a loss of a certain degree of local organization. Thus, chromosomes of LBR mutant cells have a weakened short range contact interaction compared to that of WT cells.

**Figure 5.**
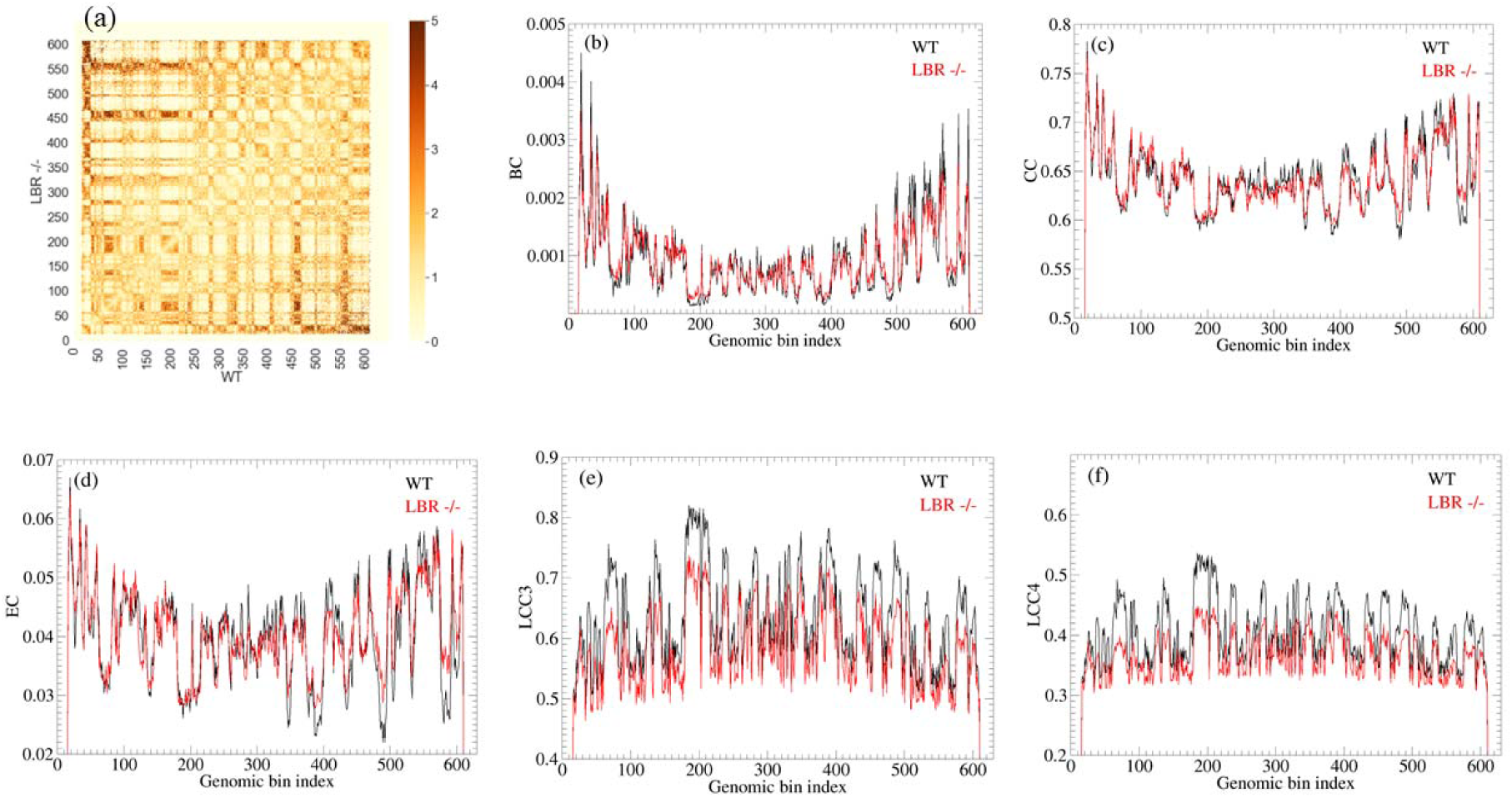
The structures of chr11 of mouse rod photoreceptor cell wildtype (WT) and LBR mutant are characterized using network properties at a resolution of 200 kb. (a) Heatmap comparison. Above the diagonal is LBR -/- cell. Below the diagonal line is WT cell. (b) BC value comparison between two the above cells. (c) CC value comparison. (d) EC value comparison. (e) LCC3 value comparison. (f) LCC4 value comparison.

As a further comparison of related cell types, we compared three cell types from different branches of the same hematopoietic lineage: erythroblasts, neutrophils and megakaryocytes. As indicated in the SM Fig. S4, the network properties of erythroblast and megakaryocyte cells are relatively close, whereas neutrophils are far from the other two. For example, the BC and LCC values for neutrophil structure have extremely high peaks around 50 bins. Such a relationship is consistent with the differentiation path of cells. Both erythroblast and megakaryocyte cells can further give rise to additional cell types (erythrocytes and platelets) in processes that involve the loss of the nucleus, while neutrophils are at a terminal stage (38). This difference may be reflected in the increased structure specialization of the neutrophil chromosomes.

### C. The connection between network properties and genomic annotations

Having observed that quantitative CSN properties differ between cell types and conditions, we next asked whether these properties of the structural network relate to other biologically significant linear annotations along the genome. A major advantage of CSN is the potential of data reduction. Various node-based network properties render the complex structure information (two-dimensional contact matrix originated from Hi-C) into different one-dimensional functions of positions (genomic bins). In this way, it is convenient to compare 1D node properties with known genomic linear annotations.

In our study, we used LAD status and chromatin states as examples to study the relationship between genomic linear annotations and network properties of the structure (LCC3, LCC4, EC, BC, and CC), focusing on IMR90 cells, as shown in Fig. 6. We also compared LAD annotation with A/B compartment strength (top eigenvector MI-PCA of Hi-C contact (40), up to a sign, Fig. 6c), where we observe that LAD labels are more likely associated with compartment B. Note that we did not fix the sign and compartment B is positive (and A is negative) for this top eigenvector of MI-PCA. Though the A/B compartment value is not a node-based network property, it nevertheless provides a value associated with each genomic position. Researchers have routinely used it to classify compartments, while more advanced methods of classification have been developed in recent years (49). As described in the Methods, we used cumulative fraction curves (CFCs) to discern the separation of LAD states (LAD versus non-LAD) by different node network properties. As shown in Fig. 6a, CFCs of LAD on EC are clearly separated from those of non-LAD and reference as well. The genomic bins associated with LAD labels are more likely to have high EC values. Similar results can also be observed for the CFC of LLC4 in Fig. 6b. In contrast, although MI-PCA can also distinguish LAD and non-LAD regions at low values, the three CFC curves representing LAD, non-LAD and reference overlap at high values (Fig. 6c).

**Figure 6.**
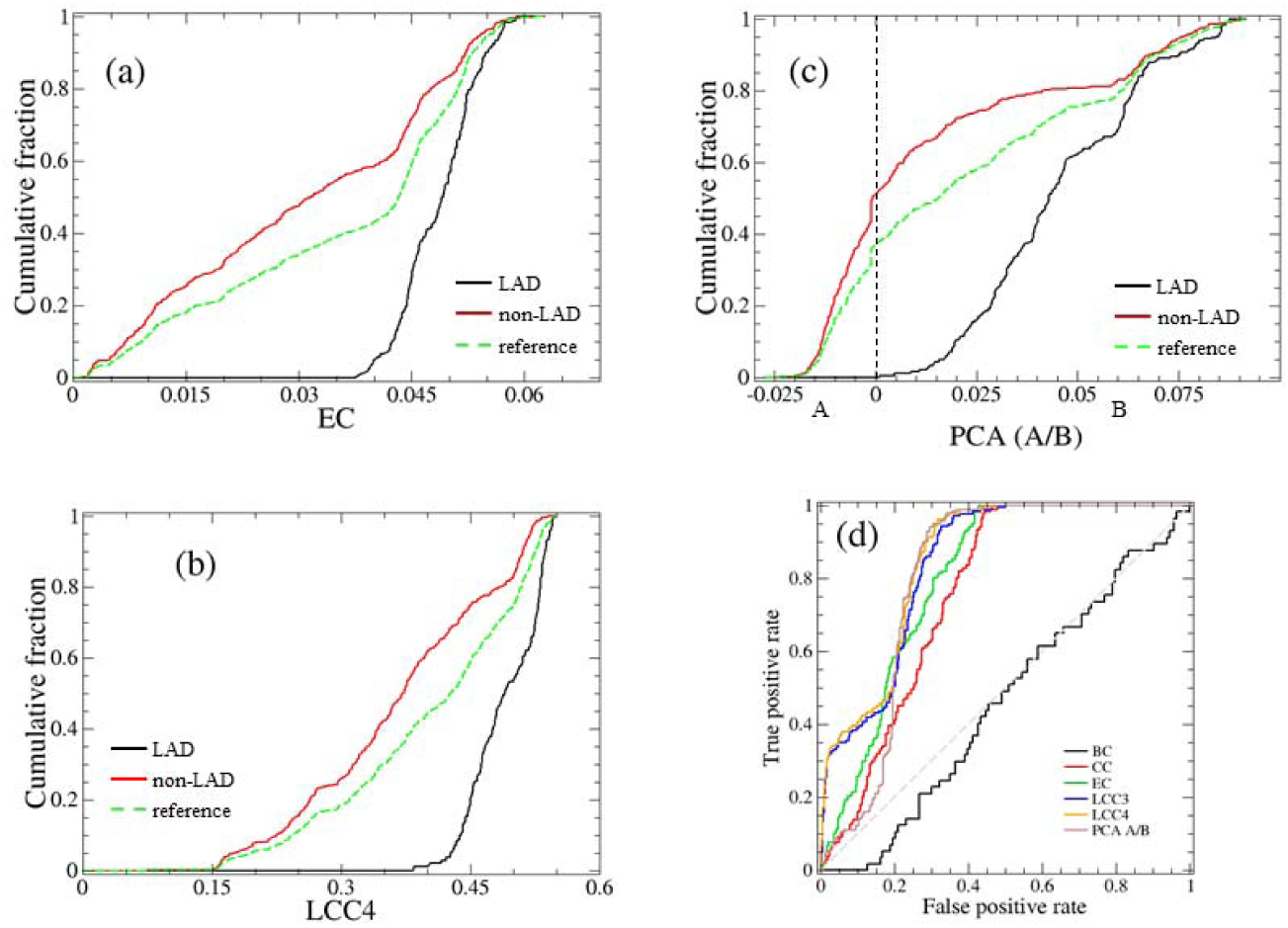
LAD cumulative fraction (CF) plots of network properties and MI-PCA for a region of IMR90 chr21. Here classifier performance concepts using CF_P_ (black), CF_N_ (red), CF_R_ (green dash) are shown for EC in (a), LCC4 in (b) and PCA in (c). (d) ROC curves of six different network properties for the LAD classification.

The comparison of how well each network property can distinguish LADs from non-LADs can be seen from the ROC graph in Fig. 6d. When an ROC is high (and thus with a larger AUC value), there is a strong separation of LAD vs. non-LAD by this metric. LCC4 values (AUC = 0.852) are the strongest predictor for a region being LAD or not, followed by EC and MI-PCA, while LAD regions show no distinction based on BC values at all. Except BC, all other five network properties are associated with differences in LAD classification. Both LCC3 and LCC4 outperform others, including the A-B compartment signal, in resolving LAD classification especially at the small false positive rate region. This suggests that LADs regions have characteristic modes of local clustering within the chromosome structure that are even more distinctive to LADs than B compartment association in general. This observation confirms the importance of quantifying different network properties to study chromosome structure and biological annotation. By examining various network properties that range from global to local, we find that LADs seem to have more distinctive local network properties rather than global.

Besides comparing LAD and MI-PCA (A/B compartment strengths), we also examined the correlation between A/B compartment subtypes and network properties. The difference between A1 and A2 sub-compartments is that A1 has higher CG content and shorter genes. B1, B2, and B3 differ in their association with histone modifications. According to the available data (45), chr21 of IMR90 cells associates with four sub-compartments A1, A2, B1, and B2. By comparing LCC3 and LCC4 vs A1, A2, B1 and B2, we found that network property LCCs can distinguish different subtypes (SM Fig. S5).

We next compared chromatin states with network properties. Chromatin states classify chromatin regions by properties such as their epigenetic marks and transcription state to identify their specific functions in transcriptional regulation (51). The chromatin states data of GM12878 chr21 (GEO:GSM936082) was used in this comparison. As shown in Fig.7a and 7b, we show cumulative curves with respect to LCC values for the six most data-rich chromatin states. Despite the similar nature of the definitions, the accumulation curves of chromatin states in LCC3 and LCC4 are distinct in discerning chromatin state classifications. LCC4 can better discriminate heterochromatin (CS13) versus other states. The distribution of either LCC metric for poised promoter state classification (CS3) is sharply peaked and there are no zero values (Fig. 7c and 7d). This result suggests that chromatin state label CS3 may have a highly specialized structure motif. In addition, some chromatin states, such as heterochromatin and repetitive/CNVs, had more than one aggregation peak at LCC4, which may indicate that there are subtypes of chromosomal structures associated with these CS labels.

**Figure 7.**
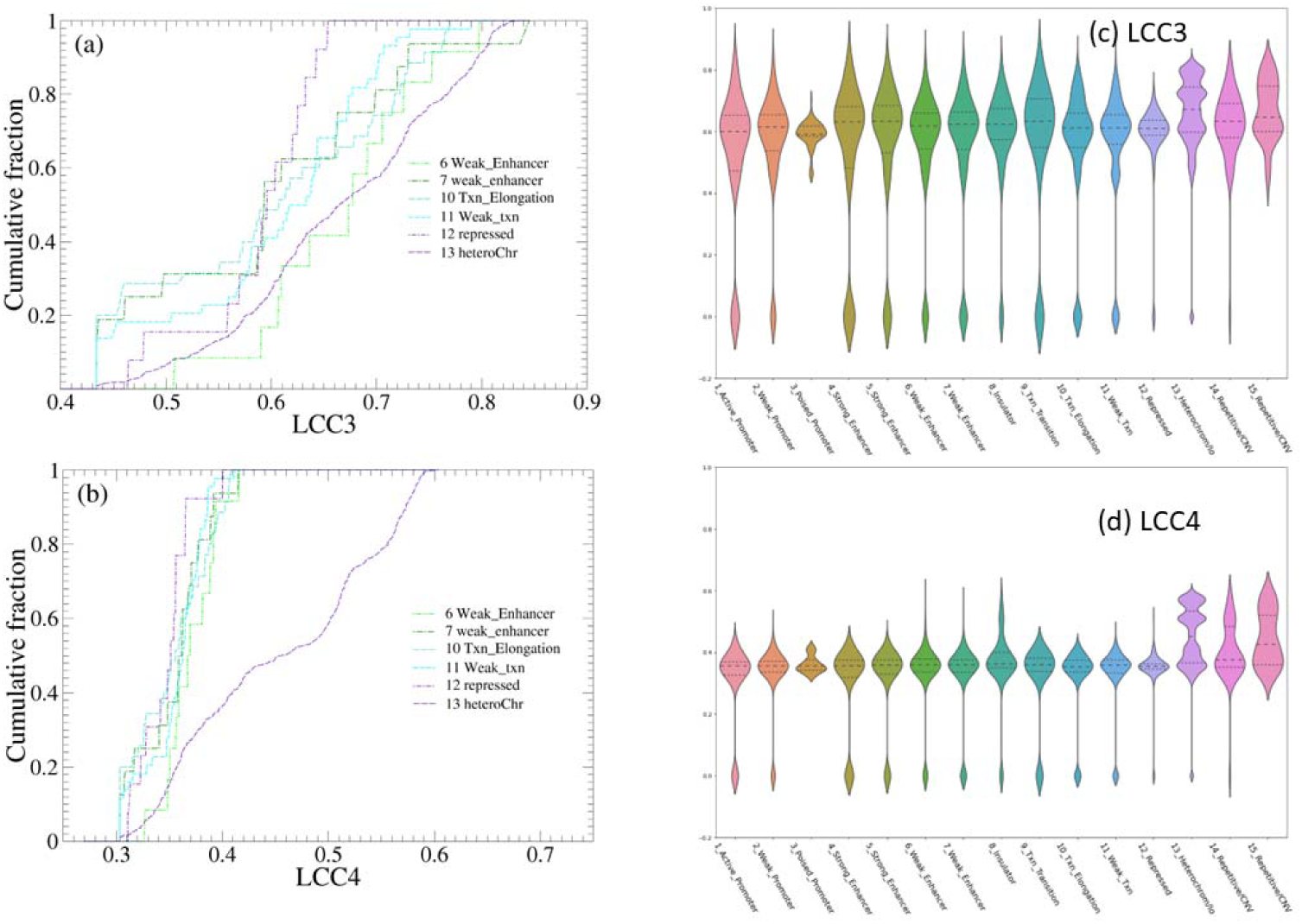
Using network properties discern selected six chromatin state labels of chr21 GM12878 for LCC3 (a) and LCC4 (b). The corresponding distribution of network properties of all 15 different chromatin states are shown in (c) and (d). The resolution is 50 kb.

## IV Concluding Remarks

We constructed the structural network of chromosomes using Hi-C sequencing data. We found that some node-based network properties, such as LCC3 and LCC4, can be used to characterize important features of chromosome structure. It can render complex two-dimensional interactions into one-dimensional quantitative network properties. Node features of CSN can be used to reveal meaning structural differences existing between cell types. For example, compared to other network properties and A/B compartment strength (contact PCA), square local clustering coefficient (LCC4) is a particularly strong classifier for predicting LAD and one specific chromatin state (CS13, heterochromatin). In general, network analysis of CSN can be a useful tool in the study of bulk Hi-C deduced three-dimensional genome structure. Similar analysis can also be used to study interchromosomal interaction and chromosome dynamics and probe other aspects of data acquisition procedures such as C-walk (52) and multi-loci interactions (53).

## Acknowledge

We acknowledge insightful discussions with D. Foutch on the network analysis of biomolecular structures.

## Notes

### Competing Interest Statement

The authors have declared no competing interest.

## References

1. Rowley, M.J. and Corces, V.G. (2018) Organizational principles of 3D genome architecture. Nat Rev Genet, 19, 789–800.

2. Bonev, B. and Cavalli, G. (2016) Organization and function of the 3D genome. Nat Rev Genet, 17, 772.

3. Robson, M.I., Ringel, A.R. and Mundlos, S. (2019) Regulatory Landscaping: How Enhancer-Promoter Communication Is Sculpted in 3D. Molecular Cell, 74, 1110–1122.

4. Liu, H., Tsai, H., Yang, M., Li, G., Bian, Q., Ding, G., Wu, D. and Dai, J. (2023) Three-dimensional genome structure and function. MedComm (2020), 4, e326.

5. da Costa-Nunes, J.A. and Noordermeer, D. (2023) TADs: Dynamic structures to create stable regulatory functions. Curr Opin Struct Biol, 81, 102622.

6. Bruckner, D.B., Chen, H., Barinov, L., Zoller, B. and Gregor, T. (2023) Stochastic motion and transcriptional dynamics of pairs of distal DNA loci on a compacted chromosome. Science, 380, 1357–1362.

7. McCord, R.P., Kaplan, N. and Giorgetti, L. (2020) Chromosome Conformation Capture and Beyond: Toward an Integrative View of Chromosome Structure and Function. Mol Cell, 77, 688–708.

8. Corsi, F., Rusch, E. and Goloborodko, A. (2023) Loop extrusion rules: the next generation. Curr Opin Genet Dev, 81, 102061.

9. Dixon, J.R., Selvaraj, S., Yue, F., Kim, A., Li, Y., Shen, Y., Hu, M., Liu, J.S. and Ren, B. (2012) Topological domains in mammalian genomes identified by analysis of chromatin interactions. Nature, 485, 376–380.

10. Crane, E., Bian, Q., McCord, R.P., Lajoie, B.R., Wheeler, B.S., Ralston, E.J., Uzawa, S., Dekker, J. and Meyer, B.J. (2015) Condensin-driven remodelling of X chromosome topology during dosage compensation. Nature, 523, 240–244.

11. Zhang, C., Xu, Z., Yang, S., Sun, G., Jia, L., Zheng, Z., Gu, Q., Tao, W., Cheng, T., Li, C. et al. (2020) tagHi-C Reveals 3D Chromatin Architecture Dynamics during Mouse Hematopoiesis. Cell Rep, 32, 108206.

12. Heinz, S., Texari, L., Hayes, M.G.B., Urbanowski, M., Chang, M.W., Givarkes, N., Rialdi, A., White, K.M., Albrecht, R.A., Pache, L. et al. (2018) Transcription Elongation Can Affect Genome 3D Structure. Cell, 174, 1522-+.

13. Belaghzal, H., Borrman, T., Stephens, A.D., Lafontaine, D.L., Venev, S.V., Weng, Z., Marko, J.F. and Dekker, J. (2021) Liquid chromatin Hi-C characterizes compartment-dependent chromatin interaction dynamics. Nat Genet, 53, 367–378.

14. García-Domenech, R., Gálvez, J., de Julián-Ortiz, J.V. and Pogliani, L. (2008) Some New Trends in Chemical Graph Theory. Chemical Reviews, 108, 1127–1169.

15. Atilgan, C., Okan, O.B. and Atilgan, A.R. (2012) In Rees, D. C. (ed.), Annual Review of Biophysics, Vol 41, Vol. 41, pp. 205–225.

16. Amitai, G., Shemesh, A., Sitbon, E., Shklar, M., Netanely, D., Venger, I. and Pietrokovski, S. (2004) Network analysis of protein structures identifies functional residues. Journal of Molecular Biology, 344, 1135–1146.

17. Di Paola, L. and Giuliani, A. (2015) Protein contact network topology: a natural language for allostery. Current Opinion in Structural Biology, 31, 43–48.

18. Doshi, U., Holliday, M.J., Eisenmesser, E.Z. and Hamelberg, D. (2016) Dynamical network of residue–residue contacts reveals coupled allosteric effects in recognition, catalysis, and mutation. Proceedings of the National Academy of Sciences, 113, 4735–4740.

19. Daily, M.D., Upadhyaya, T.J. and Gray, J.J. (2008) Contact rearrangements form coupled networks from local motions in allosteric proteins. Proteins-Structure Function and Bioinformatics, 71, 455–466.

20. Albert, R. and Barabási, A.-L. (2002) Statistical mechanics of complex networks. Reviews of Modern Physics, 74, 47–97.

21. Yao, X.Q., Momin, M. and Hamelberg, D. (2018) Elucidating Allosteric Communications in Proteins with Difference Contact Network Analysis. J Chem Inf Model, 58, 1325–1330.

22. Sethi, A., Tian, J., Vu, Dung M. and Gnanakaran, S. (2012) Identification of Minimally Interacting Modules in an Intrinsically Disordered Protein. Biophysical Journal, 103, 748–757.

23. Kryven, I., Duivenvoorden, J., Hermans, J. and Iedema, P.D. (2016) Random Graph Approach to Multifunctional Molecular Networks. Macromolecular Theory and Simulations, 25, 449–465.

24. Stauffer, D. (1985) Introduction to Percolation Theory. Taylor & Francis.

25. Zhang, B. and Horvath, S. (2005) A general framework for weighted gene co-expression network analysis. Stat Appl Genet Mol Biol, 4, Article17.

26. Hao, D., Ren, C. and Li, C. (2012) Revisiting the variation of clustering coefficient of biological networks suggests new modular structure. BMC Syst Biol, 6, 34.

27. Panditrao, G., Bhowmick, R., Meena, C. and Sarkar, R.R. (2022) Emerging landscape of molecular interaction networks: Opportunities, challenges and prospects. Journal of Biosciences, 47, 24.

28. Newman, M. (2018) Networks. OUP Oxford.

29. Solovei, I., Kreysing, M., Lanctot, C., Kosem, S., Peichl, L., Cremer, T., Guck, J. and Joffe, B. (2009) Nuclear architecture of rod photoreceptor cells adapts to vision in mammalian evolution. Cell, 137, 356–368.

30. Falk, M., Feodorova, Y., Naumova, N., Imakaev, M., Lajoie, B.R., Leonhardt, H., Joffe, B., Dekker, J., Fudenberg, G., Solovei, I. et al. (2019) Heterochromatin drives compartmentalization of inverted and conventional nuclei. Nature, 570, 395–399.

31. Javierre, B.M., Burren, O.S., Wilder, S.P., Kreuzhuber, R., Hill, S.M., Sewitz, S., Cairns, J., Wingett, S.W., Varnai, C., Thiecke, M.J. et al. (2016) Lineage-Specific Genome Architecture Links Enhancers and Non-coding Disease Variants to Target Gene Promoters. Cell, 167, 1369–1384 e1319.

32. Kaul, A., Bhattacharyya, S. and Ay, F. (2020) Identifying statistically significant chromatin contacts from Hi-C data with FitHiC2. Nat Protoc, 15, 991–1012.

33. Ay, F., Bailey, T.L. and Noble, W.S. (2014) Statistical confidence estimation for Hi-C data reveals regulatory chromatin contacts. Genome Res, 24, 999–1011.

34. Imakaev, M., Fudenberg, G., McCord, R.P., Naumova, N., Goloborodko, A., Lajoie, B.R., Dekker, J. and Mirny, L.A. (2012) Iterative correction of Hi-C data reveals hallmarks of chromosome organization. Nat Methods, 9, 999–1003.

35. Rao, S.S., Huntley, M.H., Durand, N.C., Stamenova, E.K., Bochkov, I.D., Robinson, J.T., Sanborn, A.L., Machol, I., Omer, A.D., Lander, E.S. et al. (2014) A 3D map of the human genome at kilobase resolution reveals principles of chromatin looping. Cell, 159, 1665–1680.

36. Caldarelli, G., Pastor-Satorras, R. and Vespignani, A. (2004) Structure of cycles and local ordering in complex networks. The European Physical Journal B, 38, 183–186.

37. Fronczak, A., Hołyst, J.A., Jedynak, M. and Sienkiewicz, J. (2002) Higher order clustering coefficients in Barabási–Albert networks. Physica A: Statistical Mechanics and its Applications, 316, 688–694.

38. Lind, P.G., González, M.C. and Herrmann, H.J. (2005) Cycles and clustering in bipartite networks. Physical Review E, 72, 056127.

39. Yin, H., Benson, A.R. and Leskovec, J. (2018) Higher-order clustering in networks. Physical Review E, 97, 052306.

40. Das, P., Golloshi, R., McCord, R.P. and Shen, T. (2020) Using contact statistics to characterize structure transformation of biopolymer ensembles. Physical Review E, 101, 012419.

41. Haarhuis, J.H.I., van der Weide, R.H., Blomen, V.A., Yanez-Cuna, J.O., Amendola, M., van Ruiten, M.S., Krijger, P.H.L., Teunissen, H., Medema, R.H., van Steensel, B., et al. (2017) The Cohesin Release Factor WAPL Restricts Chromatin Loop Extension. Cell, 169, 693–707 e614.

42. Kind, J., Pagie, L., de Vries, S.S., Nahidiazar, L., Dey, S.S., Bienko, M., Zhan, Y., Lajoie, B., de Graaf, C.A., Amendola, M., et al. (2015) Genome-wide Maps of Nuclear Lamina Interactions in Single Human Cells. Cell, 163, 134–147.

43. Das, P., San Martin, R. and McCord, R.P. (2023) Differential contributions of nuclear lamina association and genome compartmentalization to gene regulation. Nucleus, 14, 2197693.

44. Ernst, J., Kheradpour, P., Mikkelsen, T.S., Shoresh, N., Ward, L.D., Epstein, C.B., Zhang, X., Wang, L., Issner, R., Coyne, M. et al. (2011) Mapping and analysis of chromatin state dynamics in nine human cell types. Nature, 473, 43–49.

45. Xiong, K. and Ma, J. (2019) Revealing Hi-C subcompartments by imputing inter-chromosomal chromatin interactions. Nat Commun, 10, 5069.

46. Wen, Z., Zhang, W., Zhong, Q., Xu, J., Hou, C., Qin, Z.S. and Li, L. (2022) Extensive Chromatin Structure-Function Associations Revealed by Accurate 3D Compartmentalization Characterization. Front Cell Dev Biol, 10, 845118.

47. Nichols, M.H. and Corces, V.G. (2021) Principles of 3D compartmentalization of the human genome. Cell Rep, 35, 109330.

48. Yaffe, E. and Tanay, A. (2011) Probabilistic modeling of Hi-C contact maps eliminates systematic biases to characterize global chromosomal architecture. Nat Genet, 43, 1059–1065.

49. Kariti, H., Feld, T. and Kaplan, N. (2023) Hypothesis-driven probabilistic modelling enables a principled perspective of genomic compartments. Nucleic Acids Res, 51, 1103–1119.

50. Fatima, U., Hina, S. and Wasif, M. (2023) A novel global clustering coefficient-dependent degree centrality (GCCDC) metric for large network analysis using real-world datasets. Journal of Computational Science, 70, 102008.

51. Ernst, J. and Kellis, M. (2017) Chromatin-state discovery and genome annotation with ChromHMM. Nat Protoc, 12, 2478–2492.

52. Olivares-Chauvet, P., Mukamel, Z., Lifshitz, A., Schwartzman, O., Elkayam, N.O., Lubling, Y., Deikus, G., Sebra, R.P. and Tanay, A. (2016) Capturing pairwise and multi-way chromosomal conformations using chromosomal walks. Nature, 540, 296–300.

53. Dotson, G.A., Chen, C., Lindsly, S., Cicalo, A., Dilworth, S., Ryan, C., Jeyarajan, S., Meixner, W., Stansbury, C., Pickard, J. et al. (2022) Deciphering multi-way interactions in the human genome. Nat Commun, 13, 5498.

