## Supplementary Materials for "Node Features of Chromosome Structure Network and Their Connections to Genome Annotation"

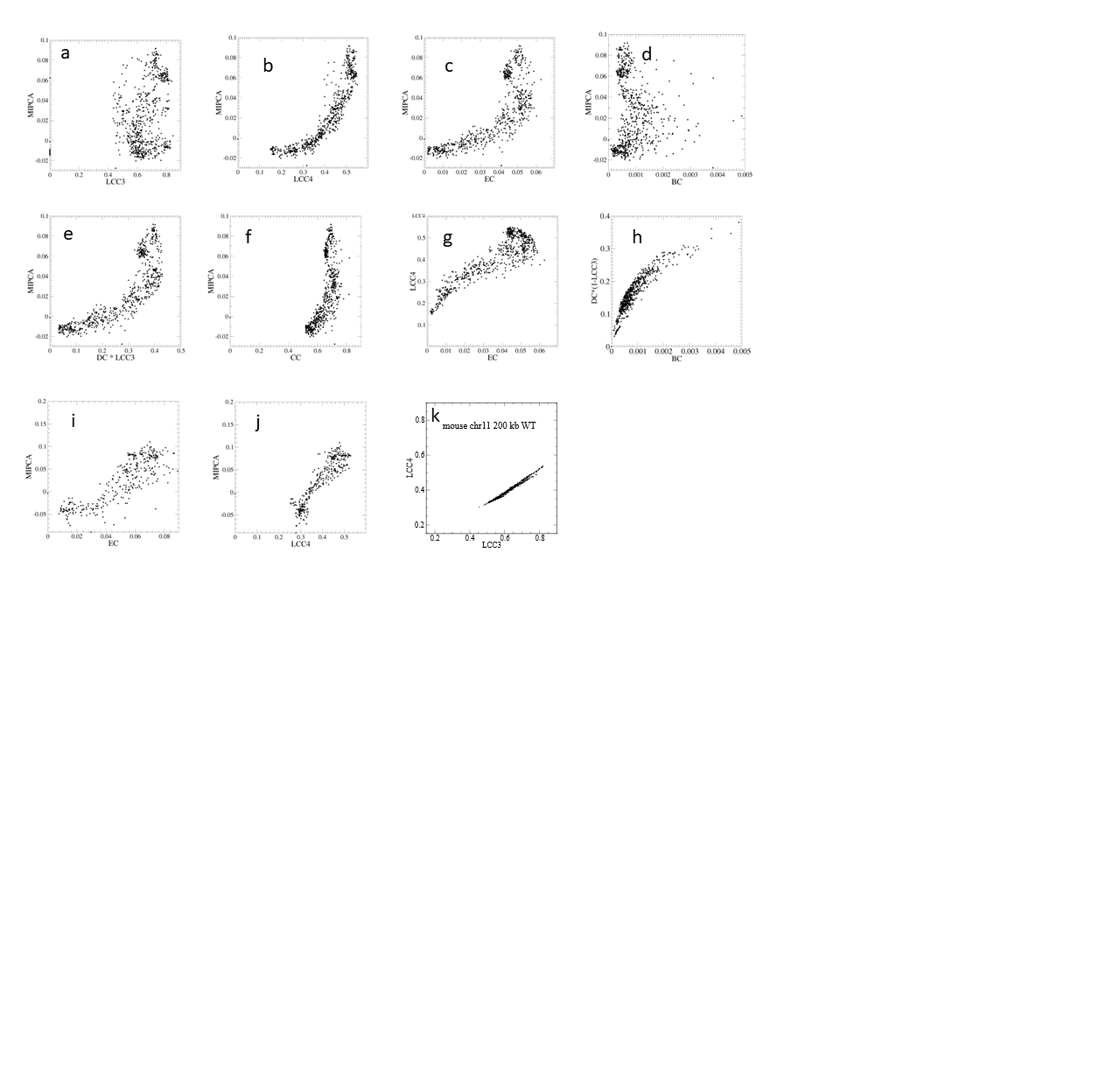

SM Figure S1: (a-f) Scatter plots of IMR90 chr21: 15.25–48.25 Mb at 50 kb resolution network properties vs. MIPCA (A/B compartment). (g-h) Correlation between different network properties using same system as above. (i-j) Network properties vs. MIPCA (A/B compartment) using IMR90 chr10: 0.0–17.3 Mb at 50 kb resolution. (k) The correlation between LCC3 and LCC4 of Mouse rod photoreceptor cells chr11 at 200 kb resolution.

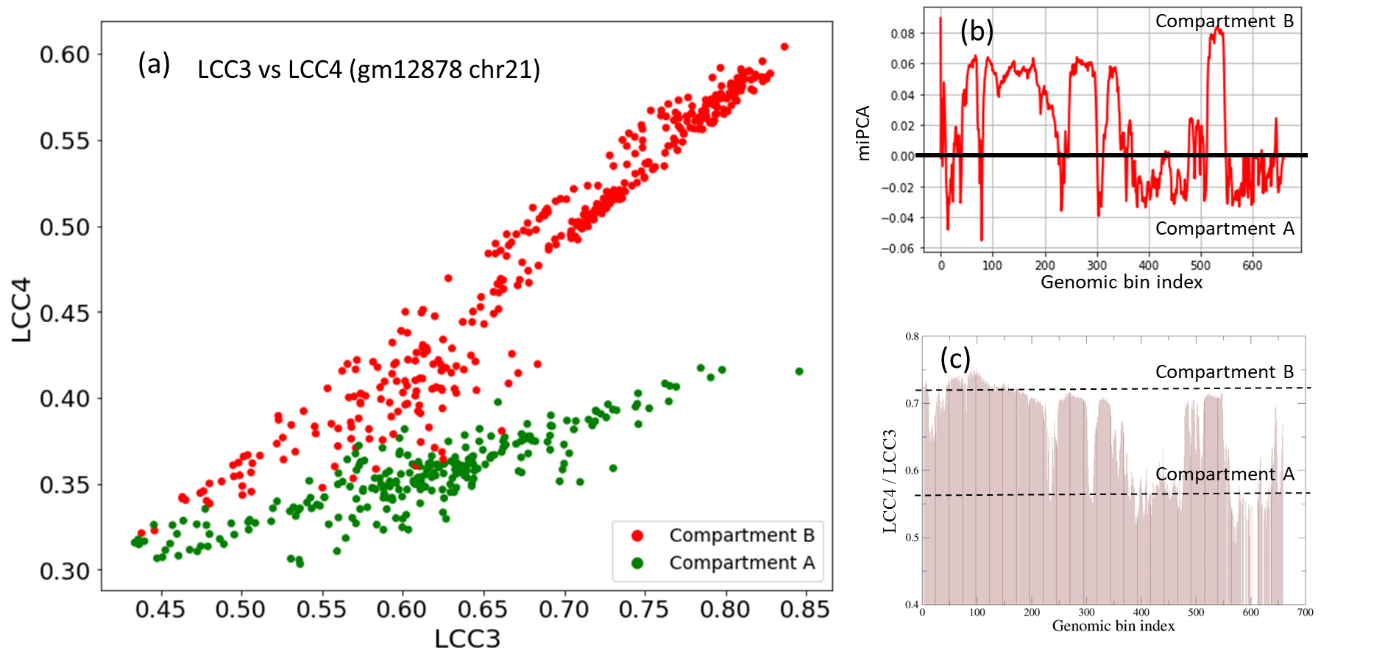

SM Figure S2: (a) Scatter plot of LCC3 and LCC4 values of GM12878 chr21. (b) GM12878 chr21 A/B compartment classified by MIPCA. (c) The ratio of LCC3/LCC4. Two slope values similar to those in panel (a) are marked with dashed lines. All above test used GM12878 chr21:15.25–48.25 Mb at 50 kb resolution.

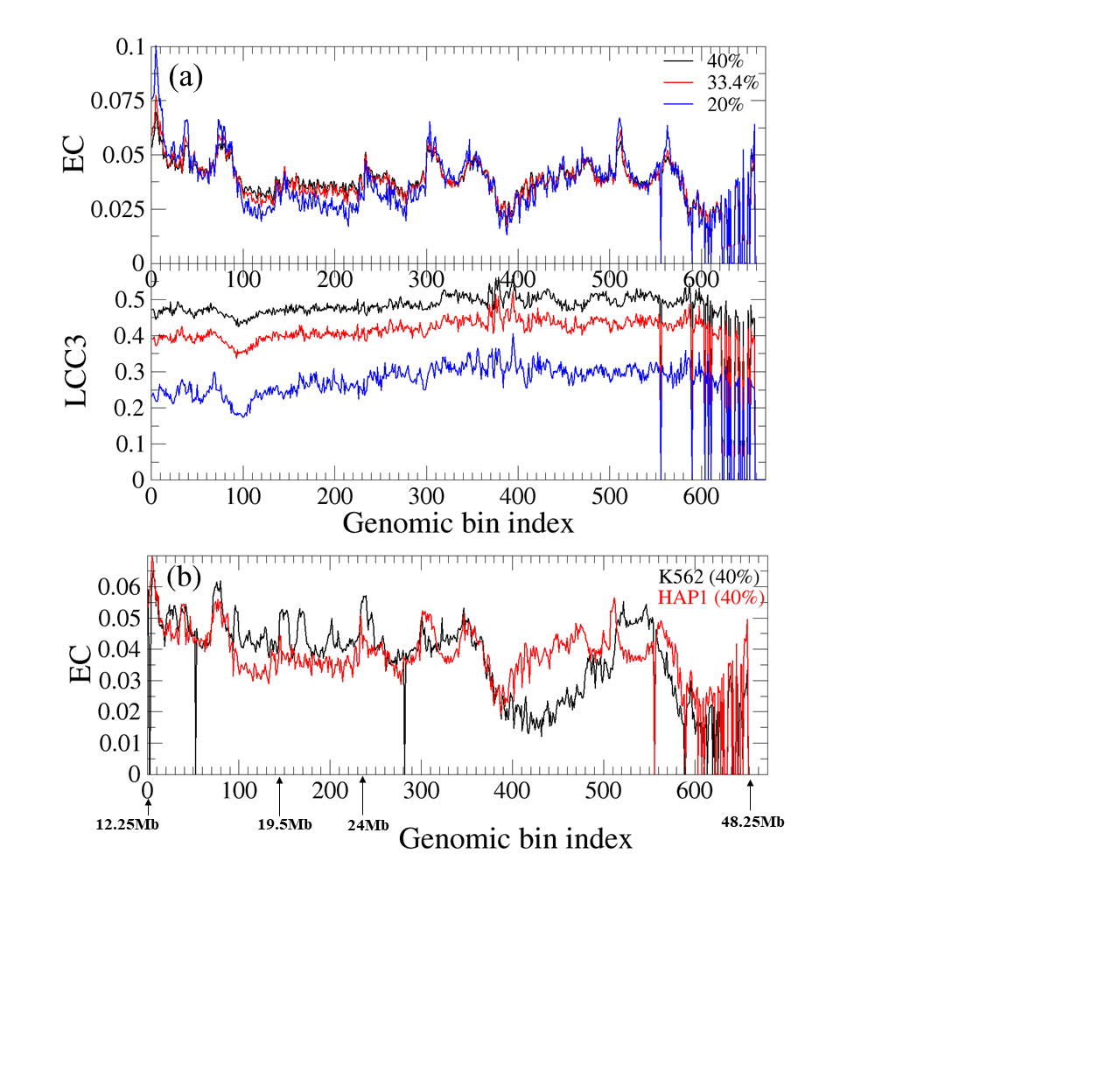

SM Figure S3: (a) Network properties with different coverage of Hap1 cells chr21: 15.25–48.25 Mb at 50 kb resolution. (b) EC as functions of genomic node index for K562 versus HAP1. Chromosome regions and resolution is same as above. Constant coverage of 40% is used.

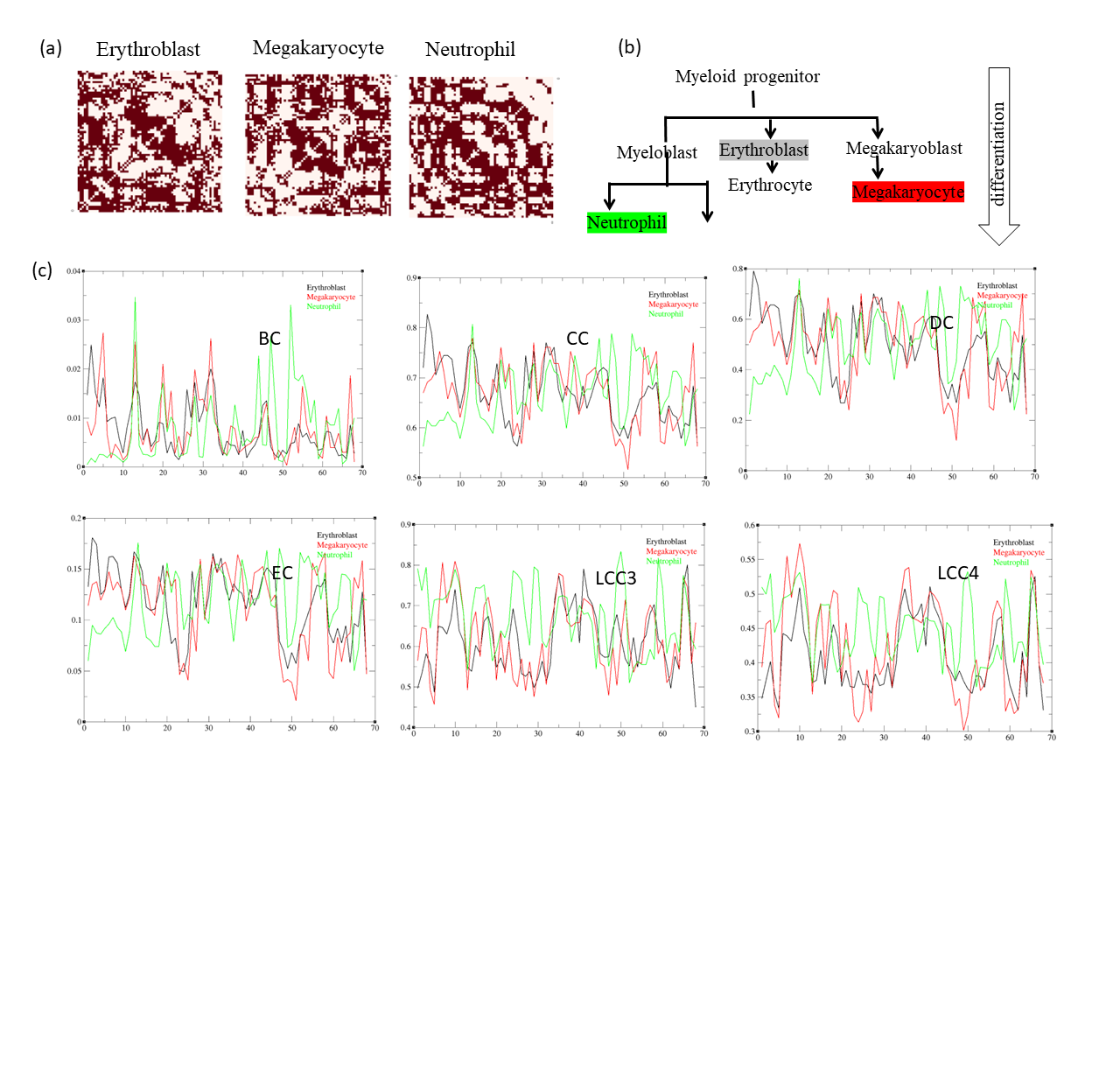
SM Figure S4: Comparison of contact networks of three different human blood cells erythroblast, megakaryocyte and neutrophil (chr10: 0–17 Mb, 250 kb resolution): (a) Comparison of the contact network pattern of the three types of cells (coverage 49%). (b) Selected blood cell lineage showing developmental relationship. (c) Comparison of network properties between these three cell types (color-coded: black, erythroblast; red, megakaryocyte; green, neutrophil).

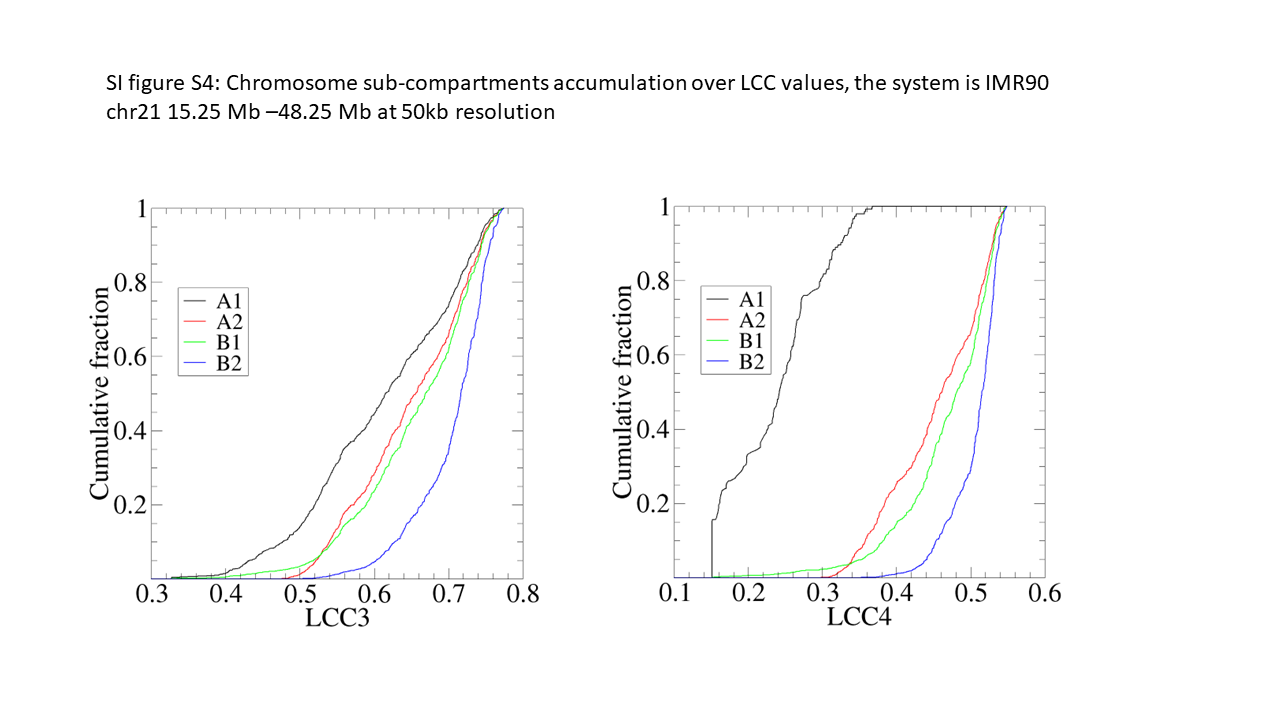
SM Figure S5: Chromosome sub-compartments accumulation over LCC values, the system is IMR90 chr21 15.25–48.25 Mb at 50 kb resolution.
